# TaHL-PTM: Post-Translational Modification Prediction in Proteins via Target-Hooked Discriminative Fine-Tuning of Decoder-only Protein Language Models

**DOI:** 10.64898/2026.08.24.746791

**Authors:** Bhawana Prasain, Pawel Pratyush, Stefan Schulze, B. KC Dukka

## Abstract

Post-translational modifications (PTMs) regulate protein function, making accurate residue-level PTM prediction essential for understanding cellular mechanisms and disease pathways. While decoder-only protein language models (PLMs) pretrained with the causal language modeling (CLM) objective have driven breakthroughs across various bioinformatics tasks, their potential for PTM prediction remains largely underexplored. CLM-based PLMs that rely on Byte-Pair Encoding (BPE) for tokenization, such as ProtGPT2, introduce intra-token label collision by merging multiple amino acids with conflicting labels into a single token, creating a major bottleneck for residue-level tasks. To overcome this, we propose TaHL-PTM (Target-Hooked Low-rank adaptation for PTM prediction), a novel framework that integrates target-hooked tokenization with site-directed discriminative LoRA fine-tuning. Target-hooked tokenization constrains tokenization around the candidate residue using dedicated marker tokens to eliminate intra-token label collision while preserving the surrounding sequence context, whereas the proposed discriminative objective repurposes the standard generative CLM objective for residue-level PTM classification by directly optimizing the separation between modified and unmodified sites. We benchmark TaHL-PTM across six distinct PTM tasks on ProtGPT2 and ProGen2 models. TaHL-PTM consistently improves MCC, with the largest gain of up to +0.11 for tyrosine phosphorylation (0.34 to 0.45), alongside improvements in F1, AUROC, and AUPR. Performance gains are more pronounced for collision-affected samples, validating the effectiveness of target-hooked tokenization, while consistent improvements across both BPE-based and per-residue-based causal PLMs demonstrate that the proposed framework generalizes across models with different pretraining tokenization schemes.

## 1 Introduction

Post-translational modifications (PTMs) are covalent chemical alterations that involve the formation or cleavage of bonds on the protein backbone or amino acid side chains [1]. These modifications occur after or during protein synthesis and regulate nearly every aspect of protein function, including stability, localization, and molecular interactions [2]. Consequently, PTMs play a central role in biological processes such as signal transduction, gene regulation, and cellular homeostasis [3]. Dysregulation of PTM mechanisms has been associated with numerous diseases, including cancer, neurodegenerative disorders, and metabolic conditions [4]. Although advances in large-scale proteomics have accelerated PTM discovery, experimental identification methods such as mass spectrometry remain expensive and are often limited by sample complexity [5]. As a result, computational prediction has become an important complementary approach for large-scale PTM discovery.

Early computational methods relied on traditional machine learning models built on handcrafted sequence features, such as hydrophobicity, charge, accessible surface area, and chain flexibility. Subsequent approaches, including MusiteDeep [6] and DeepPhos [7], improved PTM site prediction by learning sequence patterns directly from raw protein windows without manual feature engineering. These advances were further enabled by transformer-based architectures [8], which rely on self-attention mechanisms to model long-range dependencies in sequential data. More recently, protein language models (PLMs) such as ESM-2 [9], ProtBERT [10], and ProtT5 [10] have substantially improved many downstream protein analysis tasks, including protein-protein interaction prediction and PTM prediction, by learning rich contextual representations directly from protein sequences [11]. Most current PTM prediction frameworks, including recent approaches such as LMCrot [12], StackGlyEmbed [13] and UniPTMs [14], use encoder-based PLMs primarily as feature extractors, with the learned embeddings subsequently used for downstream classification.

On the other hand, relatively few studies explore the direct adaptation of large PLMs for PTM prediction instead of using them solely for feature extraction. MTPrompt-PTM [15] applies prompt tuning [16] to a structure-aware PLM [17]. Other recent work, such as PTM-Mamba [18], further investigates an alternative sequence-modeling approach based on state-space models. Despite the transformative success of causal language modeling (CLM) across generative sequence design, adapting decoder-only PLMs for residue-level PTM prediction remains underexplored. A notable exception is PTMGPT2 [19], which adapts ProtGPT2 [20] for PTM site prediction using its standard autoregressive next-token prediction objective. At the same time, PTMGPT2 also inherits the underlying tokenization strategy of ProtGPT2, raising an important question about how tokenization affects fine-grained residue-level prediction. Consequently, optimally tailoring causal decoder-only models for residue-level PTM prediction remains an open challenge.

PLMs adopt different tokenization strategies to represent amino acid sequences. Encoder-based PLMs such as ProtT5 [10] and ESM-[9], as well as decoder-based models like ProGen2 [21], operate on per-residue tokenization, where each amino acid is represented as an individual token. However, treating residues individually may limit the explicit representation of local semantic information arising from structural context and recurring sequence motifs [22]. In contrast, CLM-based decoder-only PLMs such as ProtGPT2 [20] utilize Byte-Pair Encoding (BPE), which merges frequently co-occurring amino acid subsequences into single tokens representing short motifs. By compressing the effective sequence length, BPE improves computational efficiency and enables modeling of longer protein sequences. Since the computational complexity of transformer architectures scales quadratically with sequence length [23], this compression reduces both memory usage and computational cost. Despite these advantages, BPE-based tokenization sacrifices residue-level granularity. Many protein prediction tasks, including solvent accessibility [24], protein binding site prediction [25, 26], and PTM site prediction [27, 28], require predictions at individual amino acid positions. Because a single BPE token can span multiple residues, including a mix of modified and unmodified sites, assigning a residue-level label to the entire token introduces intra-token label collision. As a consequence, BPE-based PTM models such as PTMGPT2 may receive supervision from a representation that does not uniquely correspond to the residue being classified, introducing ambiguity during training and inference.

To address this limitation, we develop TaHL-PTM (Target-Hooked Low-rank adaptation for PTM prediction), which adapts CLM-based decoder-only PLMs to residue-level PTM prediction. TaHL-PTM introduces target-hooked tokenization to explicitly isolate the target residue from multi-residue BPE tokens, thereby preventing intra-token label collisions while preserving the surrounding sequence context. Additionally, unlike PTMGPT2, which retains the conventional autoregressive objective for label-token prediction, TaHL-PTM reformulates the task as conditional classification by restricting supervision to target-site prediction and directly optimizing the PTM classification decision. Although primarily motivated by BPE-based architectures such as ProtGPT2, TaHL-PTM is equally applicable to per-residue-based decoder-only PLMs such as ProGen2. Experimental results across five PTM tasks on ProtGPT2, ProGen2-small, and ProGen2-medium demonstrate that TaHL-PTM consistently improves PTM prediction while harnessing the rich contextual representation capabilities of autoregressive PLMs.

## 2 Methods

### 2.1 Dataset Description

The training and testing datasets used in this study are derived from the benchmark curated by Wen et al. [29], which contains high-quality PTM annotations from PTMAtlas. In this benchmark, positive samples are defined as high-confidence, experimentally identified modification sites that undergo strict quality control, with a false discovery rate (FDR) *≤* 1% and a site localization probability *>* 0.5. Negative samples are defined as unmodified residues of the same amino acid type within the same proteins, for which no supporting evidence of modification is available from tandem mass spectrometry (MS/MS) experiments or the literature. We evaluate our framework across six major PTM types: phosphorylation at serine (S), threonine (T), and tyrosine (Y); arginine (R) methylation; lysine (K) acetylation; and lysine (K) SUMOylation. These PTMs represent diverse biochemical modification mechanisms and are widely adopted as standard benchmarks in PTM site prediction research [27]. The dataset is partitioned into a training set and an independent test set using a 90:10 ratio. The provided splits are constructed such that no UniProt identifiers are shared between the training and test sets, thereby preventing leakage of global protein-level information. In addition, potential homology bias and sequence redundancy between the partitions were mitigated by filtering test protein sequences against the training set using the CD-HIT algorithm [30] with a 0.9 cut-off. For a comprehensive description of the curation and preprocessing procedures for these datasets, we refer the reader to the original publication by Wen et al. [29]. The distribution of the training and testing sets is reported in Supplementary Section S1. To further assess the generalization performance of the proposed models, we additionally evaluated them on the external independent test set from Han et al. [15], which contains experimentally validated PTM annotations curated from a distinct source. Finally, for each site in the datasets, a fixed-length sequence window of 51 residues centered on the target residue is extracted and used as input to the TaHL-PTM framework.

### 2.2 Definition and Quantification of Intra-Token Label Collision

We first formalize the relationship between residue-level labels and BPE tokens. Let *r*_*t*_ denote the target residue at sequence position *t*, with label *y*_*t*_ *∈* {0, 1}. Let *r*_*i*_ denote the residue at sequence position *i*, with the corresponding label *y*_*i*_. Under BPE tokenization, the token *T*_*j*_ containing *r*_*t*_ may include multiple residues grouped together as:

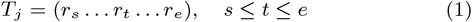

where *s* and *e* denote the start and end residue indices in the window sequence spanned by token *T*_*j*_ . Thus, the token may include residues to the left of *r*_*t*_ (*s < t*), to the right of *r*_*t*_ (*t < e*), or on both sides (*s < t < e*). Let 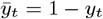 denote the complementary class label of the target residue. We define intra-token label collision as:

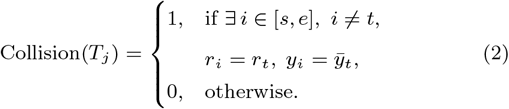

Let *N* ^train^ and *N* ^test^ denote the number of target residues in the training and test sets, respectively. The percentage of samples exhibiting intra-token label collision is computed as:

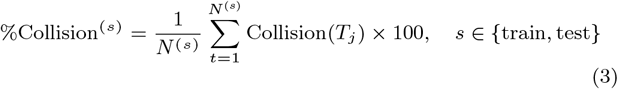

High collision rates indicate that tokenization frequently merges modified and unmodified residues, which introduces conflicting supervision during training.

### 2.3 Overview of TaHL-PTM framework

Figure 1 presents a schematic overview of the TaHL-PTM training framework. Given a protein sequence window centered on a candidate PTM residue, TaHL-PTM first applies marker-guided tokenization to obtain a target-aware token representation, which is then incorporated into structured input prompts. The prompts are subsequently processed by a causal PLM adapted via parameter-efficient Low-Rank Adaptation (LoRA), with optimization guided by the proposed Site-Directed Discriminative Fine-Tuning Objective (SDFO). The resulting decoder representation is then passed to the discriminative classification head, where it is projected to the vocabulary space to obtain class-specific logits for residue-level PTM prediction. Overall, TaHL-PTM is composed of three core components: (i) marker-guided tokenization and prompt formulation, (ii) parameter-efficient adaptation of a pretrained causal PLM, and (iii) the SDFO and discriminative training. The tokenization and prompt formulation methodology is described in Section S2.4, the implementation details of LoRA are provided in Section S2.6, and the mathematical formulation of the SDFO, including causal loss masking, the discriminative classification head, probability estimation, and the final optimization objective, is presented in Section S2.5.

**Figure 1.**
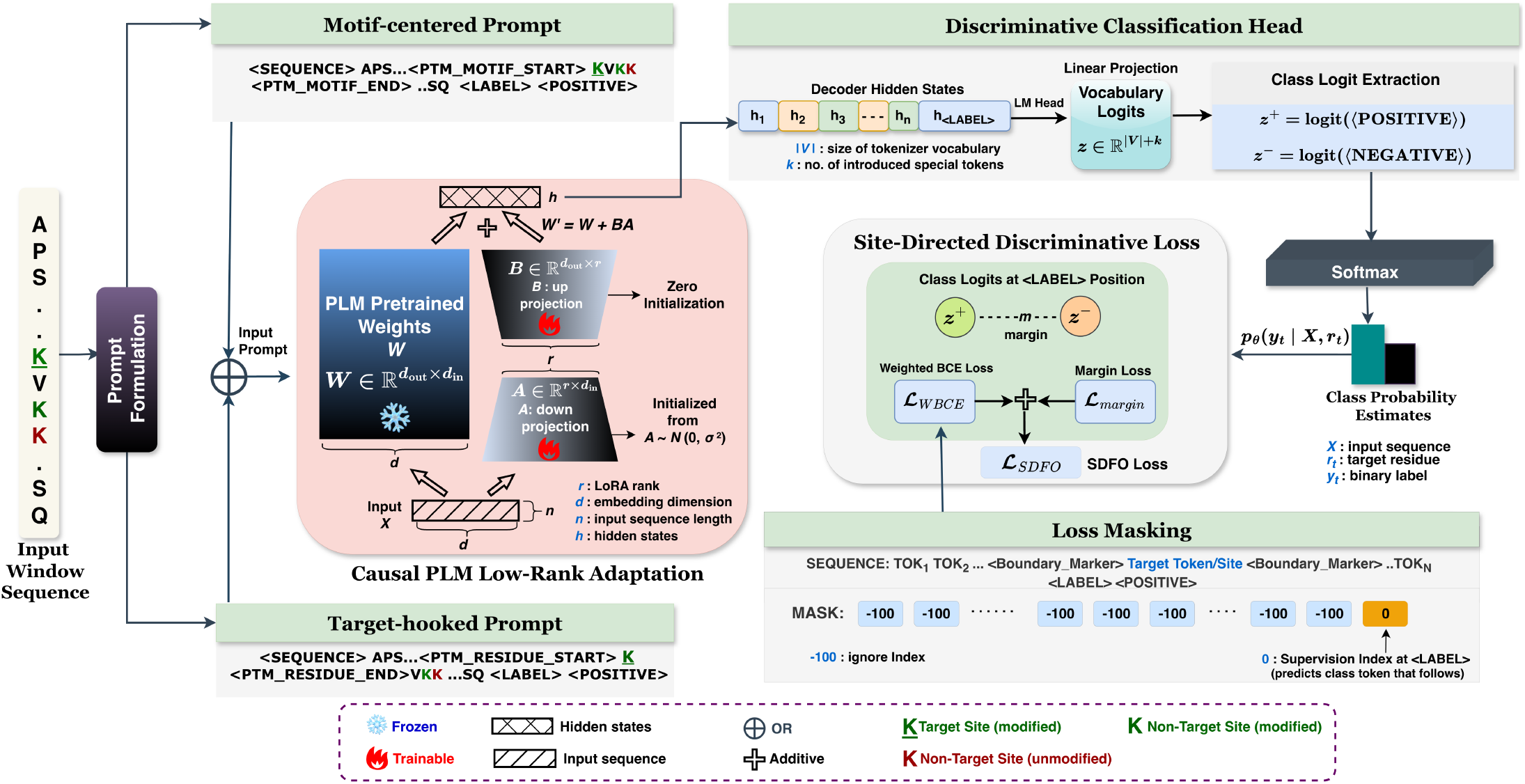
Architecture of the TaHL-PTM training framework illustrated for lysine PTM prediction. The underlined green lysine (K) denotes the target residue (modified in this example). Other non-target lysine residues within the sequence are colored green (modified) or red (unmodified), highlighting the presence of multiple nearby annotated residues that may be merged into a single BPE token.

### 2.4 Prompt Formulation

To mitigate intra-token label collisions introduced by BPE, we propose structured prompt formulations that augment the input sequence with task-specific special tokens and boundary marker tokens. Specifically, these prompts include the sequence indicator token `<SEQUENCE>`, which marks the start of the input sequence; the prediction indicator token `<LABEL>`, which designates the position from which the model autoregressively generates the class-specific tokens `<POSITIVE>` and `<NEGATIVE>`; and a set of boundary marker tokens determined by the adopted tokenization strategy. It should be noted that the class tokens `<POSITIVE>`and `<NEGATIVE>` are appended to the input prompt exclusively during training. During inference, the prompt is truncated at the `<LABEL>` token, and the model generates the class token corresponding to the predicted PTM label.

As illustrated in Figure 2 (a), a fixed 51-residue sequence window centered on the candidate residue is subjected to conventional BPE tokenization. A residue centered in sequence space is not guaranteed to be centered in token space because BPE forms tokens containing a variable number of amino acids by merging frequently co-occurring amino acid patterns. This misalignment prevents the candidate residue from being uniquely represented at the token level. Consequently, the target residue may be merged into a multi-residue token (i.e., `KVKK`), obscuring the target site and resulting in intra-token label collision. To explicitly anchor the prediction target within the token space, we introduce prompt formulations constructed using boundary-marker-guided tokenization strategies and examine two distinct design variants, as depicted in Figure 2 (b). In the motif-centered design, the boundary marker tokens *<*PTM MOTIF START*>* and *<*PTM MOTIF END*>* delimit the PTM motif while preserving the original BPE tokenization. By contrast, in the second variant, referred to as the target-hooked design, the boundary markers *<*PTM RESIDUE START*>* and *<*PTM RESIDUE END*>* are inserted immediately around the target residue prior to tokenization. The exact formulations of the two prompting strategies are provided below.

**Figure 2.**
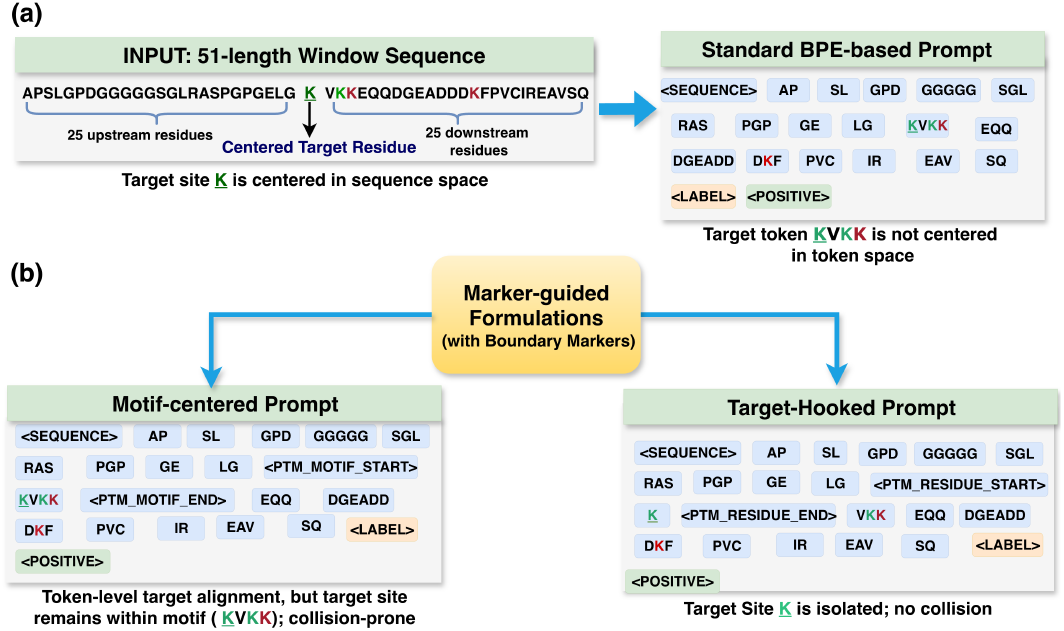
Prompt formulations in TaHL-PTM. (a) Prompt design using standard BPE tokenization, where multi-residue tokens cause intra-token label collision, (b) Marker-guided prompt formulations contrasting a motif-centered with a target-hooked tokenization-based design. Demonstrated for lysine (K) PTM prediction, modified lysines are indicated in green and unmodified lysines in red. The underlined green lysine (K) denotes the candidate target residue undergoing modification.

#### Motif-centered Design

Let *X* = (*r*_1_*r*_2_ … *r*_*n*_) denote an amino acid sequence of length *n*, where *r*_*i*_ *∈ A* and *A* denotes the amino acid alphabet. A BPE tokenizer maps the residue sequence to a sequence of BPE tokens,

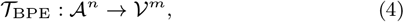

where *V* denotes the tokenizer vocabulary and *m* is the number of tokens produced for the input sequence. The resulting token sequence is:

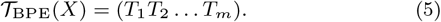

where each token *T*_*j*_ = (*r*_*s*_ … *r*_*e*_) corresponds to a contiguous subsequence of residues. Let *r*_*t*_ denote the target residue. Under standard BPE tokenization, there exists a token *T*_*j*_ such that *r*_*t*_ *∈ T*_*j*_ . The token *T*_*j*_ may therefore span the target residue together with adjacent residues. To construct the input prompt, the token sequence is partitioned with respect to *T*_*j*_, and boundary markers are inserted around it. The final prompted training input is given by:

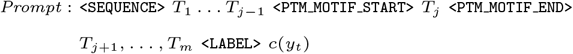

where *c*(*y*_*t*_) *∈* {`<POSITIVE>, <NEGATIVE>`} denotes the class token corresponding to the ground-truth label *y*_*t*_ and is appended to the prompt only during training. In this formulation, the boundary markers identify the BPE token *T*_*j*_ containing the target residue without altering its internal composition. As illustrated in Figure 2 (b) (left branch), we can see that in motif-centered design, the target token KVKK contains both modified and unmodified lysine residues. Thus, the motif-centered design provides token-level target alignment by identifying the token containing the prediction target, but it does not uniquely identify the target residue within that token. Therefore, intra-token label collision remains unresolved.

#### Target-Hooked Design

In this formulation, a constrained tokenization operation is proposed that isolates the target residue prior to tokenization. The *n*-length input window is decomposed as

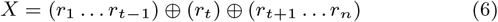

BPE is applied independently to the left and right subsequences:

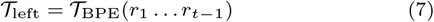

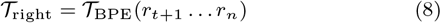

The target residue *r*_*t*_ is mapped to a singleton token 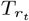, which prevents it from being merged with neighboring residues and thus avoids intra-token label collision. The resulting token sequence *T*_hooked_ is constructed as

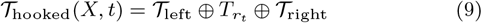

The prompted input is formed by inserting boundary markers around the isolated target token 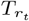 and appending special tokens. The final training input prompt is given by:

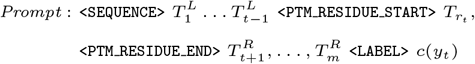

In the prompt, the superscripts *L* and *R* denote the token sequences on the left and right sequence contexts, respectively, with respect to the target residue *r*_*t*_. This operation is illustrated in Figure 2 (b) (right branch), where it isolates the target lysine residue as an independent token `K,` eliminating intra-token label collision for the prediction target. Although the neighboring token VKK in this sample still contains multiple conflicting lysine residues, they do not include the target residue. If one of these residues is selected as the target in another training instance, it is likewise isolated before tokenization.

### 2.5 SDFO: Site-Directed Discriminative Fine-Tuning Objective

To adapt CLM-based decoder-only PLMs for residue-level PTM prediction, we propose SDFO, a classification objective that reformulates autoregressive sequence modeling as a residue-level prediction task. Although these models are inherently pretrained for sequence generation, the SDFO conditions the decoders on marker-guided prompted input that explicitly identifies the prediction target. Let *V* denote the tokenizer vocabulary with size |*V*| for each PLM. The tokenizer is extended with additional special tokens (`<SEQUENCE>, <LABEL>, <POSITIVE>, <NEGATIVE>`, and marker tokens set based on the tokenization mode). Accordingly, the model’s input embedding matrix *E ∈* ℝ^|*V*|*×d*^ is expanded to ℝ^(|*V*|+*k*)*×d*^ by appending learnable embeddings corresponding to these newly introduced tokens, where *d* is the embedding dimension and *k* denotes the number of newly introduced special tokens. At the `<LABEL>` position, the autoregressive decoding process is restricted to a single prediction step. Instead of generating a sequence, the model produces a single token corresponding to the class label (`<POSITIVE>/<NEGATIVE>`). Thus, the standard next-token prediction mechanism is repurposed as a conditional classification objective, where the model estimates *p*_*θ*_(*y*_*t*_ | *X, r*_*t*_) by supervising only the logits at the `<LABEL>` position, where *X* is the input sequence, *r*_*t*_ is the target residue, *y*_*t*_ *∈* {0, 1} is its corresponding binary label, and *θ* denotes the parameters of the PLM. During training, token IDs for all positions before the `<LABEL>` position are masked (assigned an ignore index of *−*100). This prevents the model from optimizing the standard CLM objective of predicting the next sequence token, since the target sequence before `<LABEL>` corresponds to the shifted input sequence. Consequently, the gradients are only computed from the prediction of the token `<POSITIVE>` or `<NEGATIVE>` at the `<LABEL>` position. Let **h**_<LABEL>_ *∈* ℝ^*d*^ denote the hidden representation produced by the decoder at the `<LABEL>` position. This hidden representation is projected through the language modeling head to produce logits over the extended vocabulary:

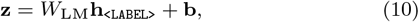

where *W*_LM_ *∈* ℝ^(|*V*|+*k*)*×d*^ and **b** *∈* ℝ^|*V*|+*k*^ denote the parameters of the language modeling head. Hence, **z** *∈* ℝ^|*V*|+*k*^ contains the logits over the extended vocabulary. From this distribution, we extract the logits corresponding to the `<POSITIVE>` and `<NEGATIVE>` tokens, denoted by *z*^+^ and *z*^*−*^, respectively. The conditional class probability estimate is obtained by applying a softmax over the two class-specific logits:

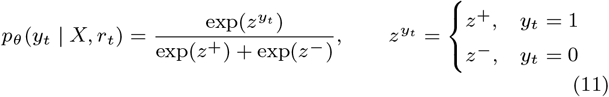

A weighted binary cross-entropy loss (*L*_WBCE_) is then optimized:

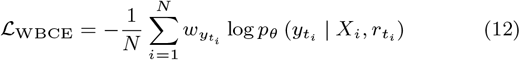

Where 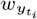 denotes the class-specific weight assigned to true label 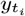 based on class frequencies to account for label imbalance across PTM datasets. To further encourage separation between positive and negative predictions, we introduce a class-weighted softplus margin loss based on the logit difference between the positive and negative class tokens. For sample *i*, let

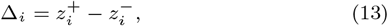

where 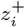 and 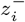 denote the logits corresponding to the `<POSITIVE>` and `<NEGATIVE>` tokens, respectively. For positive samples, the objective encourages Δ_*i*_ to be larger than a margin *m*, whereas for negative samples it encourages Δ_*i*_ to be smaller than *−m*. The margin loss is defined as

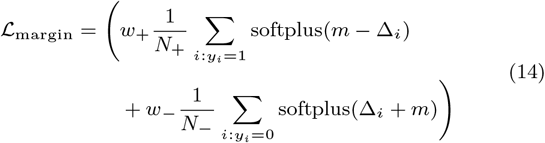

where *N*_+_ and *N*_*−*_ are the numbers of positive and negative samples in the batch, *w*_+_ and *w*_*−*_ are class-specific weights, and *m* is the margin hyperparameter. The final training loss under the SDFO is defined as:

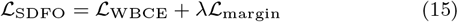

where the *λ* parameter controls the contribution of the margin term in the overall loss. The optimization objective is therefore to minimize *L*_SDFO_, jointly maximizing the conditional likelihood of the correct class while encouraging a greater separation between class-specific logits.

### 2.6 Fine-tuning Setup and Evaluation Protocol

We utilized ProtGPT2 [20] as the primary causal PLM backbone for the TaHL-PTM fine-tuning experiments, as its BPE tokenization constitutes an appropriate setting in which intra-token label collisions can arise. Fine-tuning was performed using the proposed prompting strategies under the SDFO, and unless otherwise stated, all main experiments used ProtGPT2 as the backbone. To further investigate whether our approach extends to causal PLMs with per-residue tokenization, we evaluated TaHL-PTM on two ProGen2 variants [21], ProGen2-small and ProGen2-medium. Detailed architectural specifications and tokenization properties of these PLMs are provided in Supplementary Section S2. All PLMs were fine-tuned with parameter-efficient LoRA, with adapters applied to all linear transformation layers of the decoder architecture, including the attention projection matrices for key *K*, query *Q*, and value *V*, the feed-forward input and output projections *W*_in_ and *W*_out_, and the language modeling head *W*_LM_. In addition to the LoRA parameters, the embeddings corresponding to the newly introduced special tokens were kept trainable and jointly optimized during fine-tuning. The effect of LoRA rank was evaluated over *r ∈* {4, 8, 16} with a scaling factor *α* = 2*r*. Further methodological background and implementation details on LoRA are provided in Supplementary Section S3. During training, models were optimized to minimize the SDFO loss *L*_SDFO_, using the Adam optimizer with a learning rate of 2*×*10^*−*4^, a batch size of 64, and gradient accumulation over two steps. Early stopping was applied to the validation set using the Matthews Correlation Coefficient (MCC) with a patience of 2 epochs. For the margin loss, hyperparameters *m* and *λ* were independently explored over the range [0.1, 0.5] in increments of 0.1 for each PTM task. In addition to LoRA, AdaLoRA, an adaptive-rank variant of LoRA, was evaluated as an ablation study reported in Supplementary Section S4. Hyperparameter tuning and model selection were performed using a validation set constructed from the training set, with a non-overlapping stratified validation split comprising 15% of the samples. The independent test sets were reserved exclusively for final evaluation. Model performance was assessed using four evaluation metrics: MCC, F1-score, Area Under the Receiver Operating Characteristic Curve (AUROC), and Area Under the Precision–Recall Curve (AUPR). Model reliability was further evaluated using the Expected Calibration Error (ECE), computed over ten equally spaced confidence bins. A detailed description of these evaluation metrics is provided in Supplementary Sections S7 and S9.

## 3 Experiments and results

### Target-Hooked Tokenization Improves Overall Performance, with the Largest Gains for Collision-Affected Samples

The impact of target-hooked tokenization on PTM prediction was evaluated by comparing marker-free standard BPE tokenization with marker-guided tokenization strategies. Table 1 shows that prompting based on marker-guided tokenization (i.e, motif-centered and target-hooked) provides an overall performance gains over the marker-free standard BPE approach across all PTMs. Among the marker-guided formulations, target-hooked tokenization consistently performs better than or comparably to motif-centered tokenization across the majority of PTM prediction tasks. Nonetheless, the magnitude of this improvement varies across PTM tasks, suggesting that the benefit of target-hooked tokenization may depend on the prevalence of intra-token label collision. To investigate this relationship, we quantified the prevalence of intra-token label collision using Equation (3), and the resulting statistics are reported in Table 2. As summarized in Table 2, intra-token label collision is observed in all PTM datasets, although its prevalence varies considerably.

**Table 1.** Performance comparison of marker-free, motif-centered, and target-hooked tokenization strategies across five PTM tasks on the independent test sets. Highest values are bolded in each column.

| PTM | Tokenization | MCC | F1 | AUROC | AUPR |
| --- | --- | --- | --- | --- | --- |
| Acetylation (K) | Marker-free | 0.528 | 0.779 | 0.843 | 0.818 |
|  | Motif-centered | 0.530 | 0.7783 | 0.845 | 0.830 |
|  | Target-hooked | **0.574** | **0.794** | **0.863** | **0.851** |
| SUMOylation (K) | Marker-free | 0.538 | 0.745 | 0.851 | 0.800 |
|  | Motif-centered | 0.547 | 0.751 | 0.854 | 0.803 |
|  | Target-hooked | **0.573** | **0.764** | **0.873** | **0.830** |
| Methylation (R) | Marker-free | 0.445 | 0.576 | 0.801 | 0.596 |
|  | Motif-centered | 0.445 | 0.572 | 0.811 | 0.599 |
|  | Target-hooked | **0.459** | **0.589** | **0.835** | **0.635** |
| Phosphorylation (Y) | Marker-free | 0.441 | 0.546 | 0.830 | 0.530 |
|  | Motif-centered | 0.439 | 0.546 | 0.830 | 0.530 |
|  | Target-hooked | **0.456** | **0.556** | **0.835** | **0.553** |
| Phosphorylation (ST) | Marker-free | 0.586 | 0.709 | 0.896 | 0.789 |
|  | Motif-centered | 0.590 | **0.714** | **0.898** | 0.796 |
|  | Target-hooked | **0.591** | 0.712 | **0.898** | **0.797** |

**Table 2.** Percentage of intra-token label collision in training and test sets across five PTM datasets.

| PTM | Train (%) | Test (%) | N (Train / Test) |
| --- | --- | --- | --- |
| Acetylation (K) | 3.9599 | 3.1660 | 63,300 / 5,559 |
| SUMOylation (K) | 3.9595 | 4.2948 | 71,453 / 8,615 |
| Methylation (R) | 1.5999 | 1.8206 | 49,691 / 4,449 |
| Phosphorylation (Y) | 0.2077 | 0.0343 | 58,721 / 5,814 |
| Phosphorylation (ST) | 3.4752 | 2.9214 | 612,777 / 61,202 |

If the proposed target-hooked tokenization primarily improves prediction by resolving intra-token label collision, larger performance gains should be observed for PTMs with higher collision rates. Figure 3 supports this hypothesis by comparing prediction errors for collision-affected and collision-free samples. PTMs with the highest collision rates, such as SUMOylation (4.29% test) and acetylation (3.16% test), exhibit the largest performance gains and a more pronounced reduction in prediction error for collision-affected samples under target-hooked tokenization. For example, MCC increases from 0.547 to 0.573 for SUMOylation and from 0.530 to 0.574 for acetylation, accompanied by consistent improvements in F1, AUROC, and AUPR. In contrast, methylation (1.82% test) and phosphorylation (Y) (0.03% test), which contain substantially fewer collision-affected samples, show comparatively smaller improvements. These findings indicate that the primary benefit of target-hooked tokenization arises from eliminating the contradictory supervision introduced by intra-token label collision rather than from the introduction of additional prompt tokens alone. Phosphorylation (ST) represents an interesting exception. Despite exhibiting a relatively high collision rate (2.92% in the test set), the improvement over motif-centered tokenization is comparatively modest. We attribute this to the substantially larger size of the phosphorylation (ST) training dataset (over 600,000 training samples). It should be noted that the results presented in this section are based on LoRA fine-tuning; however, a similar trend is observed under full fine-tuning (see Supplementary Section S5), suggesting that the effectiveness of target-hooked tokenization is independent of the adaptation strategy.

**Figure 3.**
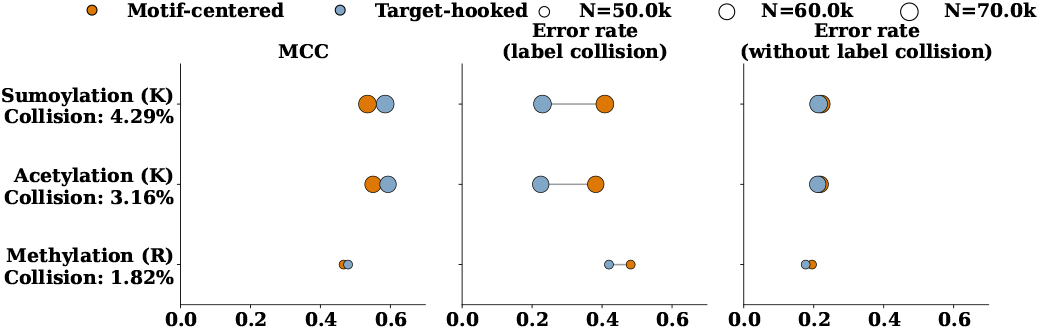
PTM-wise comparison of MCC gains and it’s association on error rates for motif centered and target-hooked tokenization on samples with and without label collison.

### TaHL-PTM Achieves Competitive Performance with Parameter-Efficient Fine-Tuning

We next examine whether LoRA-based parameter-efficient adaptation can match the performance of conventional full fine-tuning within the target-hooked TaHL-PTM framework. Across ProtGPT2, ProGen2-medium, and ProGen2-small, the relative effectiveness of the two adaptation strategies varies with both the underlying PLM and PTM task. As summarized in Table 3, LoRA achieves higher MCC than full fine-tuning for four of the five PTM tasks with ProtGPT2, with the largest improvement observed for phosphorylation (ST), where MCC increases from 0.550 to 0.591. For ProGen2-medium and ProGen2-small, the comparison is more task-dependent. LoRA achieves higher MCC for acetylation (K), SUMOylation (K), and phosphorylation (Y), whereas full fine-tuning performs better for methylation (R) and phosphorylation (ST). The largest gains from LoRA over full fine-tuning for the ProGen2 models are observed for phosphorylation (Y), with MCC improvements of +0.074 for ProGen2-medium and +0.092 for ProGen2-small. Across the three PLMs, ProtGPT2 generally shows the strongest overall performance, achieving the highest MCC in four of the five PTM tasks under full fine-tuning and three of the five tasks under LoRA. Nevertheless, the ProGen2 models show task-specific advantages, with ProGen2-small achieving the highest LoRA performance for acetylation and ProGen2-medium for phosphorylation (Y). Taken together, these results indicate two key observations: first, SDFO-guided discriminative LoRA adaptation incorporated with target-hooked tokenization remains effective across both BPE-based ProtGPT2 and per-residue ProGen2 models, indicating transferability across causal PLMs with different pretraining tokenization techniques; second, LoRA achieves performance comparable to, and in several cases better than, full fine-tuning while requiring substantially fewer trainable parameters (see Supplementary Section S3 for trainable parameter comparisons).

**Table 3.**
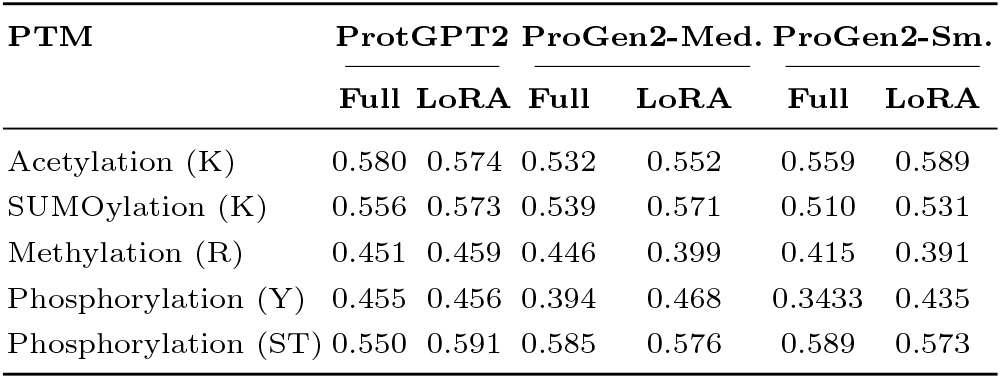
Comparison of full fine-tuning and LoRA-based parameter-efficient fine-tuning across three causal PLMs on the independent test sets. MCC is reported.

### TaHL-PTM Outperforms Existing Decoder-only PTM Prediction Method

To assess the effectiveness of TaHL-PTM, we compare it against the PTMGPT2[19] approach, retrained and evaluated on the same training and independent test sets used in this study to ensure a fair comparison. PTMGPT2 predicts label tokens using the autoregressive objective of ProtGPT2 [20] and fine-tunes all ProtGPT2 parameters. As shown in Figure 4, TaHL-PTM consistently outperforms the PTMGPT2 baseline across all five PTM prediction tasks and evaluation metrics, with improvements observed for every PTM task. The largest performance gains are observed for phosphorylation (Y) and methylation (R). Importantly, these performance improvements are achieved while updating only 0.6%– 1.6% of the model parameters, corresponding to a reduction of approximately 98.4%–99.4% in trainable parameters (depending on the selected LoRA rank; see Supplementary Section S3) compared with full fine-tuning. TaHL-PTM also achieved the highest overall performance on the additional independent benchmark from Han et al. [15] across all five PTM tasks, further supporting its cross-dataset generalization. The corresponding results are reported in Supplementary Section S8.

**Figure 4.**
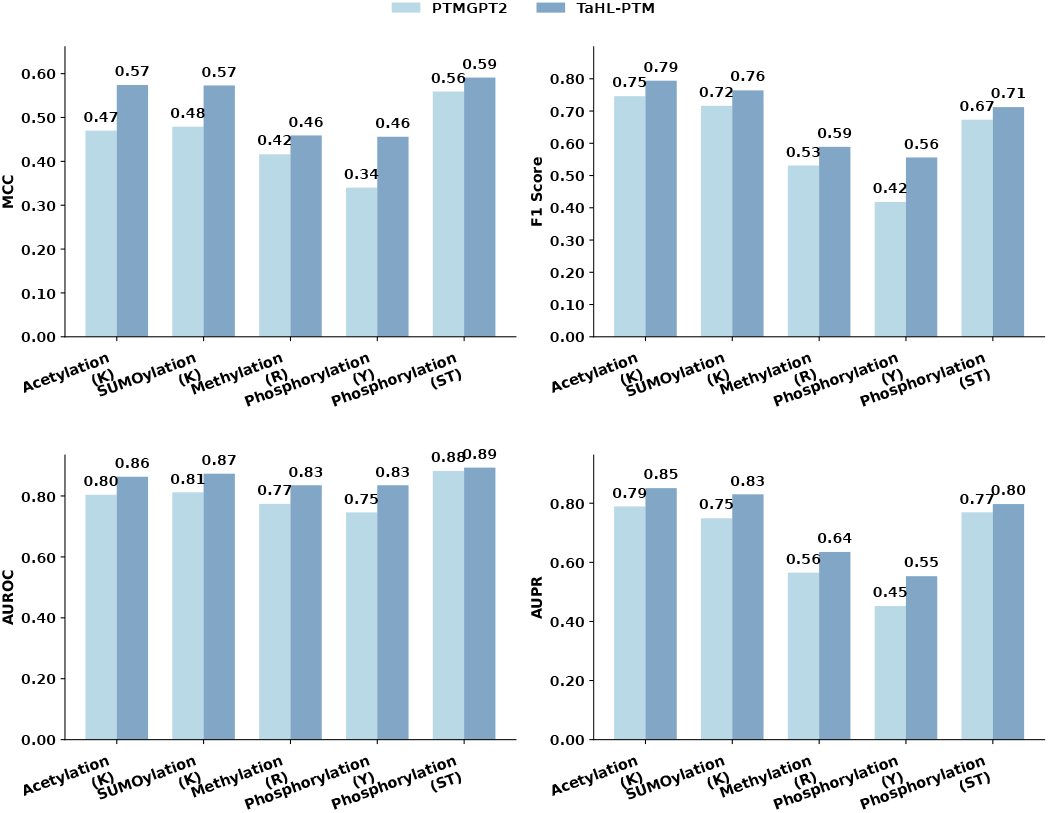
Performance comparison between PTMGPT2 and TaHL-PTM on the independent test sets across five PTM prediction tasks. The upper panel reports MCC and F1 score, while the lower panel reports AUROC and AUPR. TaHL-PTM consistently outperforms PTMGPT2 across all evaluation metrics and PTM tasks.

### Target-hooked tokenization reduces expected calibration error

Intra-token label collision introduces conflicting supervision signals, which may cause models to produce overconfident predictions despite incorrect classifications. If target-hooked tokenization effectively resolves this ambiguity, it should not only enhance predictive performance but also yield improved calibration of predicted probabilities. Therefore, we investigate whether resolving such collisions leads to better-calibrated probability estimates.

In Figure 5, we compare the calibration of three TaHL-PTM configurations: full fine-tuning with target-hooked tokenization, LoRA fine-tuning with motif-centered tokenization, and LoRA fine-tuning with target-hooked tokenization. Each bubble represents a confidence bin *b* (=10), with its x- and y-coordinates corresponding to the mean predicted confidence, confidence(*b*), and the empirical accuracy, accuracy(*b*), respectively. The size of each bubble is proportional to the number of samples in the bin, |*b*|. Perfect calibration occurs when accuracy(*b*) = confidence(*b*), corresponding to points lying on the diagonal. Across the evaluated PTM tasks, the full fine-tuned configuration generally shows larger deviations from the diagonal, indicating poorer calibration. In contrast, the LoRA-based configurations tend to cluster more closely around the diagonal, resulting in lower expected calibration error (ECE) values. Among the three configurations, target-hooked LoRA consistently achieves the lowest ECE across all PTM tasks, indicating better agreement between predicted confidence and empirical accuracy than motif-centered LoRA and thus yielding more reliably calibrated probability estimates.

**Figure 5.**
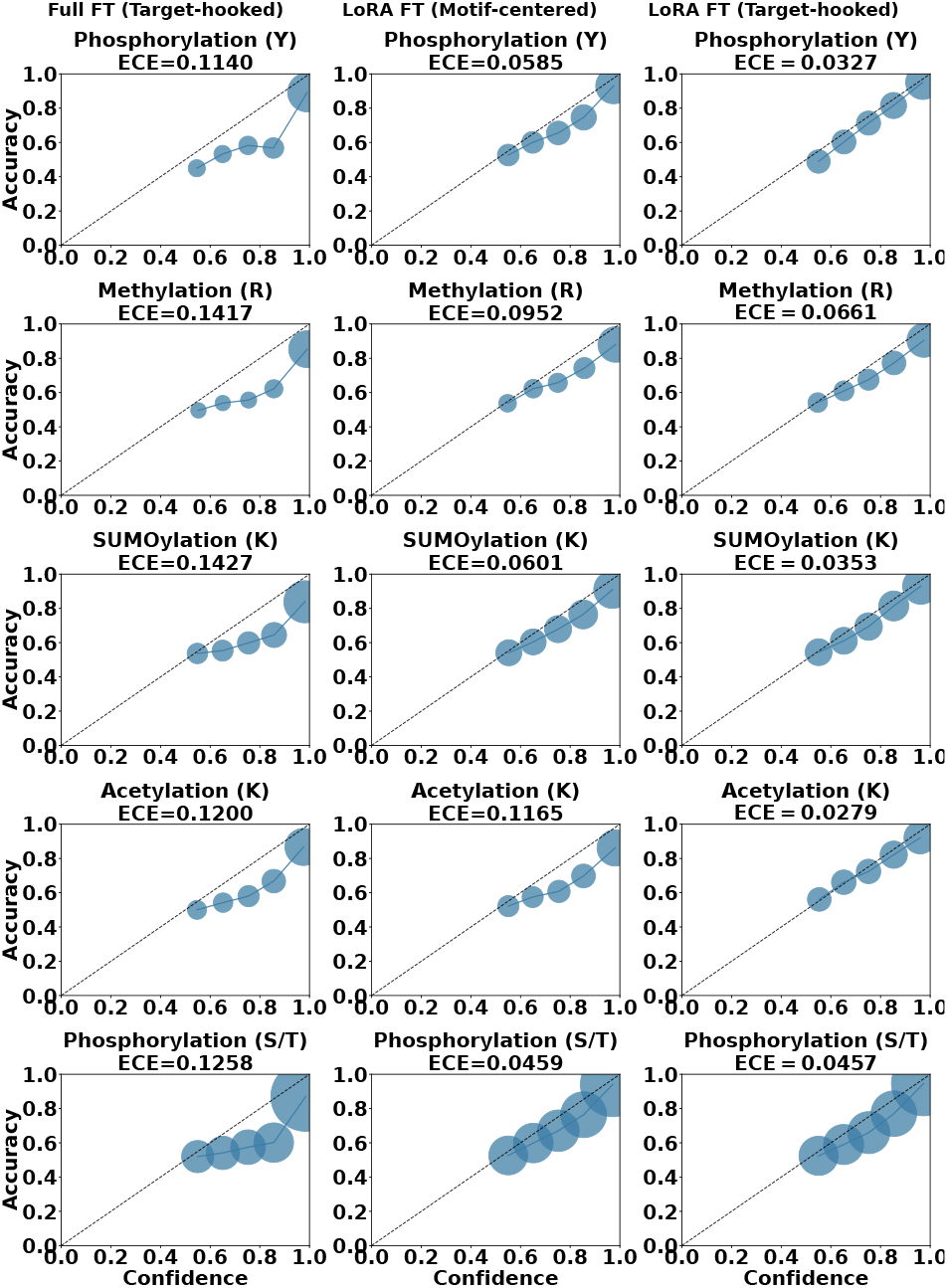
Probability calibration and Expected Calibration Error (ECE) across full and LoRA fine-tuning with motif-centered and target-hooked tokenization for five PTM prediction tasks. The diagonal represents perfect calibration.

### Target-hooked tokenization sharpens attribution at the target residue and preserves local sequence context

To better understand how the two tokenization strategies influence the sequence information used for prediction, we compared their gradient-based attribution profiles across residue positions relative to the target residue. Higher attribution scores indicate greater sensitivity of the model prediction to perturbations in the representation associated with a given residue position, reflecting the relative contribution of that position to the predicted PTM class. Across all PTM tasks, target-hooked tokenization produced a more pronounced attribution peak at the target residue (position 0), suggesting that the model assigns greater importance to the target residue position during prediction, as shown in Figure 6. This effect is particularly apparent for acetylation and SUMOylation, where attribution is strongly concentrated at the central residue. For methylation, the difference between the two tokenization strategies was less evident, although target-hooked tokenization still showed a stronger central attribution signal. This weaker distinction in the attribution profiles is consistent with the smaller performance gains observed for methylation compared with acetylation and SUMOylation.

**Figure 6.**
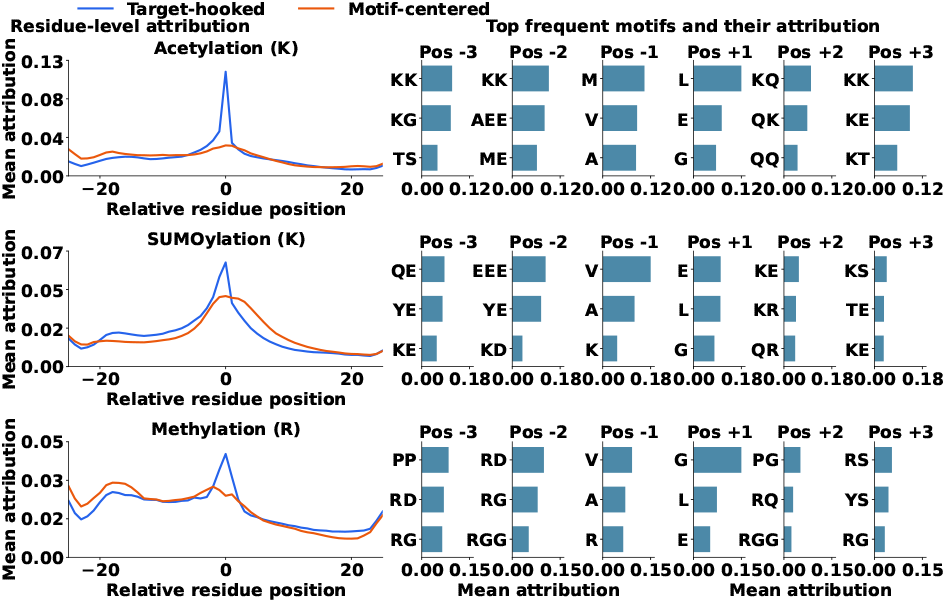
Gradient attribution profiles learned by target-hooked and motif-centered tokenization strategies. The left panels show mean attribution scores across sequence positions relative to the target residue, while the right panels display the most highly attributed motifs at neighboring positions. As shown in the left panel, target-hooked tokenization produces a stronger attribution peak at the target residue.

The right panels of Figure 6 indicate that the neighboring positions also retain substantial attribution, indicating that isolating the target residue does not prevent the model from utilizing local sequence context. For acetylation (K), attribution is concentrated on lysine-containing motifs such as KK, KQ, and KE. For SUMOylation (K), the most highly attributed neighboring motifs are enriched in lysine- and glutamate-containing sequence patterns, including KE, QE, YE, and EEE. For methylation (R), arginine-rich motifs, including RG, RGG, and RD, are the most highly attributed neighboring motifs around the site of interest. These analyses suggest that target-hooked tokenization concentrates attribution more strongly at the target residue. At the same time, appreciable attribution is retained across the flanking residues. This is consistent with the neighboring-position and motif patterns described above, indicating that the model preserves and continues to utilize local sequence context for prediction. This interpretation is also supported by the largely preserved BPE token-length distribution under target-hooked tokenization (Supplementary Figure S1), suggesting that target isolation does not substantially alter the surrounding token structure.

## 4 Conclusion

In this work, we introduced TaHL-PTM, a residue-aware parameter-efficient adaptation framework for CLM-based decoder-only PLMs. By combining target-hooked tokenization with SDFO-guided LoRA fine-tuning, TaHL-PTM enables robust residue-level PTM prediction while preserving the intrinsic modeling capabilities of CLM-based decoder-only PLMs. Across a diverse set of PTM prediction tasks and PLM backbones, our framework consistently improved predictive performance, calibration, and interpretability while requiring only a small fraction of the trainable parameters of full fine-tuning. Crucially, the most pronounced gains occur on collision-affected sequences, demonstrating the effectiveness of target-hooked tokenization in eliminating subword label ambiguity introduced by BPE tokenization. These findings establish TaHL-PTM as a powerful, generalizable, and parameter-efficient paradigm for adapting CLM-based decoder-only PLMs to residue-level protein prediction tasks, regardless of their underlying tokenization schemes.

## Supporting information

Supplementary Section

## Availability and implementation

TaHL-PTM will be publicly available soon.

## Author contributions

B.P. (Conceptualization [lead], Methodology [lead], Investigation [equal], Writing & Editing [lead]), P.P. (Conceptualization [supporting], Methodology [supporting], Investigation [equal], Writing & Editing [lead]), S.S. (Project administration [equal], Supervision [supporting], Writing & Editing [supporting]), and

D.B.K. (Conceptualization [equal], Funding acquisition [lead], Project administration [lead], Supervision [lead], Writing & Editing [supporting]).

## Acknowledgments

The authors are grateful to Dr. Christopher Homan (Rochester Institute of Technology, USA) and Dr. Meenal Chaudhari (Illinois State University, USA) for their valuable suggestions. This work was supported in part by the National Science Foundation (NSF) [NSF: #1901793 and #1564606].

