## Supplementary Section for "TaHL-PTM: Post-Translational Modification Prediction in Proteins via Target-Hooked Discriminative Fine-Tuning of Decoder-only Protein Language Models"

### Supplementary Information

### S1. Data Distribution

**Table S1** Distribution of positive and negative samples in the training and test datasets for each PTM.

| PTM | Training | | Test | |
|---|---|---|---|---|
| | Positive | Negative | Positive | Negative |
| Acetylation (K) | 30,053 | 33,247 | 2,836 | 2,723 |
| SUMOylation (K) | 35,245 | 36,208 | 3,690 | 4,925 |
| Methylation (R) | 8,616 | 41,075 | 978 | 3471 |
| Phosphorylation (S/T) | 164,879 | 447,898 | 17,184 | 44,018 |
| Phosphorylation (Y) | 10,394 | 48,327 | 1,022 | 4,792 |

Table S1 shows the number of positive and negative samples in the training and independent test sets for each PTM dataset. Dataset sizes vary across PTMs, with phosphorylation (S/T) being the largest dataset and methylation (R) the smallest. acetylation (K) and SUMOylation (K) are relatively balanced, methylation (R), phosphorylation (S/T), and phosphorylation (Y) have notable class imbalance, with more negative than positive samples.

### S2. Architectural and Tokenization Differences Across Evaluated PLMs

**Table S2** Architectural characteristics of the PLMs used in this study.

| Property | ProtGPT2 | ProGen2-small | ProGen2-medium |
|---|---|---|---|
| Parameters | 738M | 151M | 764M |
| Transformer Layers | 36 | 12 | 27 |
| Hidden Size | 1280 | 1024 | 1536 |
| Attention Heads | 20 | 16 | 16 |
| Context Length | 1024 | 1024 | 1024 |
| Vocabulary Size | 50,257 | 32 | 32 |
| Tokenization | BPE | Per-residue | Per-residue |
| Pre-training Data | UniRef50 | UniRef90+BFD30 | UniRef90+BFD30 |
| Training Objective | CLM | CLM | CLM |

We evaluated the proposed TaHL-PTM framework using two CLM based decoder-only PLMs families: ProtGPT2 [1], ProGen2[2]. For ProGen2, we experiment on small and medium version. All models are pre-trained using a causal language modeling (CLM) objective, where the model predicts the next token in a protein sequence given the preceding context. Below, we provide a detailed description of the three PLMs.

#### ProtGPT2

ProtGPT2 [1] is a GPT-2-based protein language model containing 738 million parameters. The model was trained on the UniRef50 database and consists of 36 transformer decoder layers with a hidden size of 1280 and 20 self-attention heads. ProtGPT2 supports sequence lengths up to 1024 tokens and uses Byte Pair Encoding (BPE) tokenization with a vocabulary of 50,257 tokens. Unlike per-residue tokenization schemes, BPE merges frequently co-occurring amino-acid subsequences into motif-level tokens, reducing sequence length while potentially obscuring residue-level supervision.

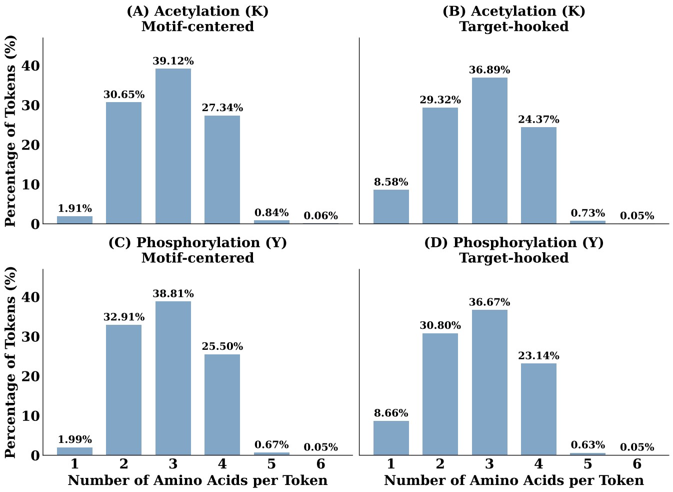


**Figure S1** Residue composition of ProtGPT2 after tokenization under motif-centered and target-hooked tokenization. Bars represent the percentage of token occurrences containing one to six amino acid residues for Acetylation (K) and Phosphorylation (Y). Target-hooked tokenization selectively increases the proportion of single-residue tokens while preserving the overall BPE token-length distribution.

For ProtGPT2, we additionally analyzed the number of amino acid residues represented by each token under both motif-centered and target-hooked tokenization strategies. Figure S1 shows the percentage of token occurrences containing one to six amino acid residues for Acetylation (K) and Phosphorylation (Y) datasets. Although tokens containing more than six amino acid residues exist, they are extremely infrequent and are not included in the figure for visual clarity. Across both datasets, tokens containing three amino acid residues are the most common, followed by tokens containing two and four residues. Target-hooked tokenization increased the proportion of single-residue tokens due to the isolated target residue introduced by the proposed strategy. Apart from this expected increase, the overall residue-per-token distribution remained largely unchanged.

#### ProGen2

ProGen2 [2] is a family of autoregressive transformer decoders trained on large-scale protein sequence databases using a next-token prediction objective. It has 4 models with different parameter sizes; ProGen2-small, ProGen2-medium, ProGen2-large and ProGen2-xlarge. The standard ProGen2 models are pre-trained on a mixture of UniRef90 and BFD30 protein sequences. The architecture follows a causal transformer decoder with rotary positional encodings. Unlike ProtGPT2, ProGen2 uses per-residue tokenization with a vocabulary size of 32, that includes 20 amino-acid and special tokens, preserving a one-to-one correspondence between residues and input tokens. Table S2 summarizes the key architectural characteristics of the evaluated PLMs.

### S3. LoRA Implementation Details and Trainable Parameter Analysis

LoRA (Low-Rank Adaptation) is a PEFT method that adapts a pretrained weight matrix $W \in \mathbb{R}^{d_{\text{out}} \times d_{\text{in}}}$ by adding a trainable low-rank update while keeping the original pretrained weights frozen. The adapted weight matrix $W'$ is defined as

$$W' = W + \Delta W = W + \frac{\alpha}{r} BA$$

where $A \in \mathbb{R}^{r \times d_{\text{in}}}$ and $B \in \mathbb{R}^{d_{\text{out}} \times r}$ are trainable low-rank matrices, $r$ is the LoRA rank, and $\alpha$ is the scaling factor. Following the LoRA initialization, $A$ is initialized from normal distribution and $B$ is initialized to zero. This initialization ensures that the model initially preserves the pretrained behavior.
In this work, the pretrained parameters of ProtGPT2 and ProGen2 models were frozen, and LoRA adapters were inserted into all linear transformation layers of the decoder, including the self-attention projections, feed-forward layers, and language modeling head. We evaluated ranks $r \in \{4, 8, 16\}$ with scaling factor $\alpha = 2r$ and LoRA dropout of 0.1. Only the LoRA adapter parameters and newly introduced special-token embeddings were updated during training of TaHL-PTM.

**Table S3** Number of trainable parameters for full fine-tuning and LoRA fine-tuning with different ranks.

| Model | Full FT | LoRA (r=4) | LoRA (r=8) | LoRA (r=16) |
|---|---|---|---|---|
| ProtGPT2 | 774,039,040 | 3,379,200 | 6,328,320 | 12,226,560 |
| ProGen2-small | 151,120,933 | 885,760 | 1,672,192 | 3,245,056 |
| ProGen2-medium | 764, 762, 149 | 2,918,400 | 5,572,608 | 10,881,024 |

The trainable parameter count for full fine-tuning is slightly larger than the original pretrained model because TaHL-PTM extends the tokenizer with additional special tokens. Consequently, the corresponding rows are appended to the input embedding matrix and the output prediction matrix (LM head), and these newly introduced parameters are optimized during fine-tuning.
Table S3 shows that compared to full fine-tuning, LoRA reduces the number of trainable parameters by approximately 98.4%–99.4% across all models and ranks. For example, ProtGPT2 has 774M trainable parameters for full fine-tuning but only 12.2M parameters with LoRA ($r = 16$), representing a 98.4% reduction. Similar reduction in parameters are observed for ProGen2-small and ProGen2-medium. This shows the high parameter efficiency of LoRA for adapting large PLMs.

### S4. Ablation on Parameter-Efficient Fine-Tuning Methods

We compare LoRA and AdaLoRA across five PTM types using three PLMs and multiple rank–alpha configurations (Figure S2). Following the original AdaLoRA formulation, the initial rank was progressively reduced to half of its original value while training using a scheduled pruning strategy. Both initialization and pruning phases accounted for 10% of the total training steps. AdaLoRA dynamically reallocates rank during training by pruning less important singular directions and assigning budget to more important ones. This introduces rank scheduling based on initial and final rank, importance estimation and pruning operations. As a result, optimization becomes more complicated. Additional hyperparameters were set to $\beta_1 = \beta_2 = 0.85$, $\Delta T = 10$, and orthogonal regularization weight of 0.5. Across different settings, there is no clear winner between LoRA and AdaLoRA.
For ProtGPT2 [1], the performance of LoRA [3] and AdaLoRA [4] is mostly comparable. In contrast, AdaLoRA shows noticeable degradation especially with ProGen2-small with rank of 8 and alpha 16. In particular, increasing the rank does not significantly improve AdaLoRA performance, suggesting that adaptive rank allocation does not effectively capture task-relevant directions in this setting.

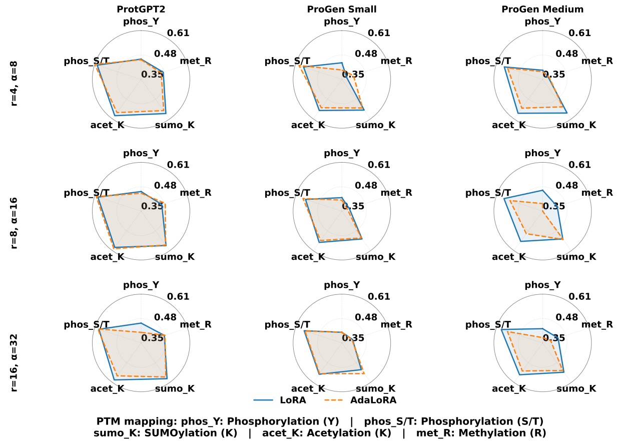


**Figure S2** Comparison of LoRA and AdaLoRA across five PTM types using three PLMs. Radar plots show MCC scores under different rank–alpha configurations. LoRA consistently achieves stronger and more stable performance across models and PTM types, while AdaLoRA exhibits variability and degradation in several settings, particularly for ProGen2-based models.

Overall, LoRA demonstrated greater consistency across rank configurations, whereas AdaLoRA showed higher variability depending on the model and task. These results suggest that the adaptive rank allocation strategy of AdaLoRA does not provide a clear advantage for PTM prediction and that fixed-rank LoRA offers a more stable and reliable parameter-efficient fine-tuning approach.

### S5. Effect of tokenization strategy on full fine-tuning

As shown in Table S4, target-hooked tokenization consistently improves prediction performance across most PTM tasks, while maintaining comparable performance for Phosphorylation (ST) under full fine-tuning. The similar trends observed under both full fine-tuning and LoRA fine-tuning demonstrate that the effectiveness of target-hooked tokenization is robust to the choice of adaptation strategy.

**Table S4.** Performance of TaHL-PTM with full fine-tuning of ProtGPT2 across five PTM types on the independent test set .

| PTM | Tokenization | MCC | F1 | AUROC | AUPR |
|---|---|---|---|---|---|
| Acetylation(K) | Marker-free | 0.522 | 0.780 | 0.838 | 0.821 |
| | Motif centered | 0.526 | 0.763 | 0.833 | 0.828 |
| | Target hooked | **0.580** | **0.806** | **0.868** | **0.855** |
| SUMOylation(K) | Marker-free | 0.520 | 0.731 | 0.846 | 0.796 |
| | Motif centered | 0.521 | 0.733 | 0.840 | 0.785 |
| | Target hooked | **0.556** | **0.757** | **0.857** | **0.807** |
| Methylation(R) | Marker-free | 0.423 | 0.557 | 0.801 | 0.578 |
| | Motif centered | 0.435 | 0.560 | 0.792 | 0.591 |
| | Target hooked | **0.451** | **0.586** | **0.815** | **0.606** |
| Phosphorylation(Y) | Marker-free | 0.427 | 0.523 | 0.828 | 0.549 |
| | Motif centered | 0.439 | 0.535 | 0.835 | 0.556 |
| | Target hooked | **0.455** | **0.550** | **0.842** | **0.572** |
| Phosphorylation(ST) | Marker-free | 0.536 | 0.677 | 0.867 | 0.756 |
| | Motif centered | 0.537 | 0.678 | **0.878** | **0.761** |
| | Target hooked | **0.550** | **0.684** | 0.875 | 0.758 |

### S6. Effect of rank in independent test set performance

**Table S5.** Evaluation on the independent test set using ProtGPT2 shows that predictive performance remains largely stable across LoRA rank values of 4, 8, and 16, except for phosphorylation (Y), indicating limited sensitivity to adaptation rank for the majority of PTM tasks.

| PTM | Rank | MCC | F1 | AUROC | AUPR |
|---|---|---|---|---|---|
| Acetylation (K) | 4 | 0.574 | 0.796 | 0.860 | 0.843 |
| | 8 | 0.572 | 0.797 | 0.858 | 0.840 |
| | 16 | 0.574 | 0.795 | 0.862 | 0.851 |
| SUMOylation (K) | 4 | 0.571 | 0.761 | 0.868 | 0.825 |
| | 8 | 0.572 | 0.766 | 0.869 | 0.823 |
| | 16 | 0.573 | 0.764 | 0.873 | 0.830 |
| Methylation (R) | 4 | 0.448 | 0.580 | 0.824 | 0.621 |
| | 8 | 0.456 | 0.586 | 0.829 | 0.624 |
| | 16 | 0.459 | 0.589 | 0.835 | 0.635 |
| Phosphorylation (Y) | 4 | 0.448 | 0.552 | 0.833 | 0.560 |
| | 8 | 0.455 | 0.556 | 0.834 | 0.553 |
| | 16 | 0.427 | 0.537 | 0.833 | 0.546 |
| Phosphorylation (ST) | 4 | 0.590 | 0.712 | 0.898 | 0.797 |
| | 8 | 0.586 | 0.709 | 0.895 | 0.792 |
| | 16 | 0.580 | 0.705 | 0.893 | 0.781 |

We evaluated multiple LoRA configurations by varying rank and corresponding scaling factor. The performance for acetylation (K), Sumoylation(K), methylation (R), and phosphorylation (ST) resulted only in marginal changes in MCC, F1, AUROC, and AUPR on increasing rank from 4 to 16. Only for phosphorylation (Y) preformance degraded on increasing rank from 8 to 16. The low sensitivity to rank for majority of PTMs suggests that adaptation capacity is not the primary bottleneck for PTM prediction in pretrained PLMs. Instead, factors such as input representation, tokenization strategy, and supervision quality are likely to have a greater impact on downstream performance than further increasing the number of trainable low-rank parameters.

### S7. Quantification of Expected Calibration Error

We calculate Expected Calibration Error(ECE) by partitioning predictions into $B = 10$ confidence bins over the interval $[0, 1]$. For each bin $b$, where $|b|$ denotes the number of samples assigned to bin $b$, we compute the empirical accuracy, accuracy$(b) = \frac{1}{|b|}\sum_{i\in b}\mathbf{1}(\hat{y}_i = y_i)$, where $\hat{y}_i$ and $y_i$ denote the predicted and true labels of sample $i$, respectively, and $\mathbf{1}(\cdot)$ is the indicator function. The mean confidence is computed as confidence$(b) = \frac{1}{|b|}\sum_{i\in b}\hat{p}_i$, where $\hat{p}_i$ is the predicted probability assigned to the predicted class for sample $i$. Calibration error is summarized using the expected calibration error (ECE), defined as $\text{ECE} = \sum_{b=1}^{B}\frac{|b|}{N}\,|\text{accuracy}(b) - \text{confidence}(b)|$, where $N$ is the total number of evaluation samples.

### S8. Cross-dataset evaluation confirms robust generalization of TaHL-PTM

| PTM | Method | MCC | F1 | AUROC | AUPR |
|---|---|---|---|---|---|
| Acetylation (K) | TaHL-PTM | **0.300** | **0.705** | **0.682** | **0.644** |
| | PTMGPT2 | 0.268 | 0.696 | 0.674 | 0.633 |
| | DeepMVP | 0.290 | 0.698 | 0.674 | 0.627 |
| SUMOylation (K) | TaHL-PTM | **0.418** | **0.740** | **0.778** | **0.756** |
| | PTMGPT2 | 0.407 | **0.740** | 0.733 | 0.688 |
| | DeepMVP | 0.380 | 0.730 | 0.753 | 0.724 |
| Methylation (R) | TaHL-PTM | **0.470** | **0.733** | **0.769** | 0.721 |
| | PTMGPT2 | 0.438 | 0.697 | 0.728 | 0.692 |
| | DeepMVP | 0.325 | 0.716 | 0.750 | **0.732** |
| Phosphorylation (Y) | TaHL-PTM | **0.390** | **0.716** | **0.747** | **0.707** |
| | PTMGPT2 | 0.303 | 0.641 | 0.707 | 0.683 |
| | DeepMVP | 0.343 | 0.702 | 0.720 | 0.692 |
| Phosphorylation (ST) | TaHL-PTM | **0.454** | **0.750** | **0.782** | **0.746** |
| | PTMGPT2 | 0.207 | 0.516 | 0.647 | 0.604 |
| | DeepMVP | 0.424 | 0.743 | 0.776 | 0.744 |

**Table S6.** Comparison of TaHL-PTM with PTMGPT2 and DeepMVP across five PTM types. The best result for each PTM and metric is shown in bold.

To evaluate the generalization capability of the proposed framework beyond the training distribution, we performed cross-dataset evaluation using the MTPrompt benchmark dataset by Han et. al. [5]. In this setting, all models were trained on the DeepMVP training set curated by Wen et al. [6] and evaluated on the independent MTPrompt dataset without any additional fine-tuning. Importantly, from the MTPrompt test dataset, all samples with overlapping UniProtID with the training set of DeepMVP are removed to prevent data leakage. We compared TaHL-PTM on this cross-dataset setting with PTMGPT2[7] and DeepMVP, both re-implemented using the same training and test sets. For DeepMVP, which combines the predictions of the ten best-performing models, we follow its inference protocol by removing outlier predictions and averaging the remaining predictions. Table S6 summarizes the comparative results across acetylation (K), SUMOylation (K), methylation (R), phosphorylation (Y), and phosphorylation (S/T).

Across all five PTM tasks, TaHL-PTM demonstrates consistent improvement in cross-dataset robustness compared with PTMGPT2, while showing superior or competitive performance compared with DeepMVP.

### S9. Evaluation Metrics

The performance of the proposed model is evaluated using four widely adopted metrics: Matthews Correlation Coefficient (MCC), F1-score, Area Under the Receiver Operating Characteristic Curve (AUROC), and Area Under the Precision–Recall Curve (AUPR). These metrics provide complementary perspectives on classification performance and are particularly suitable for evaluating PTM prediction tasks with imbalanced class distributions.

#### *S9.0.1. Matthews Correlation Coefficient (MCC)*

The Matthews Correlation Coefficient (MCC) is a balanced metric that takes into account all four outcomes of the confusion matrix: true positives (TP), true negatives (TN), false positives (FP), and false negatives (FN). It is defined as

$$\text{MCC} = \frac{TP \times TN - FP \times FN}{\sqrt{(TP + FP)(TP + FN)(TN + FP)(TN + FN)}}. \qquad \text{(S1)}$$

The MCC ranges from $-1$ to 1, where 1 indicates perfect prediction, 0 corresponds to random prediction, and $-1$ indicates complete disagreement between predictions and true labels.

#### *S9.0.2. F1-score*

The F1-score is the harmonic mean of precision and recall and provides a balanced measure of predictive performance. It is computed as

$$\text{F1} = \frac{2 \times \text{Precision} \times \text{Recall}}{\text{Precision} + \text{Recall}}, \qquad \text{(S2)}$$

where

$$\text{Precision} = \frac{TP}{TP + FP}, \qquad \text{Recall} = \frac{TP}{TP + FN}. \qquad \text{(S3)}$$

The F1-score ranges from 0 to 1, with higher values indicating better classification performance.

#### *S9.0.3. Area Under the ROC Curve (AUROC)*

The Area Under the Receiver Operating Characteristic Curve (AUROC) evaluates a model's ability to distinguish between positive and negative samples across different decision thresholds. The ROC curve is constructed by plotting the True Positive Rate (TPR) against the False Positive Rate (FPR), defined as

$$\text{TPR} = \frac{TP}{TP + FN}, \qquad \text{FPR} = \frac{FP}{FP + TN}. \qquad \text{(S4)}$$

AUROC values range from 0 to 1, where 1 represents perfect discrimination, 0.5 corresponds to random guessing, and values below 0.5 indicate worse-than-random performance.

#### *S9.0.4. Area Under the Precision–Recall Curve (AUPR)*

The Area Under the Precision–Recall Curve (AUPR) summarizes the trade-off between precision and recall across different classification thresholds. It is calculated as the area under the precision–recall curve and is particularly informative for imbalanced datasets, where the positive class is relatively rare.

The AUPR score ranges from 0 to 1, with higher values indicating superior predictive performance and a better balance between precision and recall.
